# Central platelet-rich plasma administration promotes weight loss in diet-induced obese mice

**DOI:** 10.64898/2026.10.09.758107

**Authors:** Xiaoyan Yang, Xuefang Hu, Xiaohua Yang, Qi Xu, Marcos David Munoz, Mahilini Muthamilan, Dharshini Jayaprakash, Pingwen Xu, Pei Luo

**Author notes:** To whom correspondence should be addressed: Pingwen Xu, Ph.D., 909 S. Wolcott Avenue, 2051, Chicago, IL 60612, Pei Luo, Ph.D., 909 S. Wolcott Avenue, 2040, Chicago, IL 60612. Co-first author (Xiaoyan Yang, Xuefang Hu).

## Abstract

**Aims:** Platelet-rich plasma (PRP) contains diverse bioactive factors, but its potential role in central regulation of energy balance remains unexplored. This study investigated whether central PRP administration alters metabolic homeostasis in diet-induced obese (DIO) mice and examined associated hypothalamic responses under obese and inflammatory conditions.

**Materials and Methods:** PRP was administered intracerebroventricularly in mouse models of DIO and lipopolysaccharide (LPS)-induced systemic inflammation. Body weight, food intake, energy metabolism, and adiposity were assessed alongside hypothalamic microglial morphology and transcriptomic profiling.

**Results:** A single intracerebroventricular PRP injection reduced body weight in DIO mice, with the effect persisting throughout the 7-day observation period. PRP also decreased cumulative food intake, relative gonadal white adipose tissue weight, and fasting glucose, accompanied by increased energy expenditure, oxygen consumption, fat utilization, and physical activity. In both DIO and LPS-treated mice, PRP reduced IBA1-positive area per microglial cell in the arcuate nucleus without altering microglial number. Hypothalamic RNA sequencing revealed changes in immune and metabolic signaling, extracellular matrix organization, and growth-related pathways, suggesting a complex transcriptional response rather than uniform suppression of inflammation.

**Conclusions:** A single central PRP administration induces sustained weight loss and favorable metabolic changes in DIO mice, accompanied by region-specific hypothalamic microglial and transcriptional responses. These findings identify central PRP as a potential modulator of energy balance and provide a basis for investigating its active components and underlying mechanisms.

## Introduction

Obesity reflects dysregulation of both peripheral metabolism and the central neuroendocrine mechanisms that govern energy balance. The hypothalamus coordinates feeding, energy expenditure, and glucose homeostasis by integrating hormonal and nutrient-related signals. Within the hypothalamus, the arcuate nucleus (ARC) contains neuronal populations responsive to leptin, insulin, and nutrients and distributes this information through circuits controlling whole-body metabolism^1–5^. Importantly, this metabolic integration occurs within a cellular microenvironment shaped by glia as well as neurons. Microglia, the resident innate immune cells of the central nervous system, respond to nutritional and inflammatory cues by altering their morphology and function. High-fat diet feeding and obesity induce hypothalamic inflammatory and glial responses^6–8^, and accumulating evidence implicates ARC microglia in diet-associated neuronal dysfunction and metabolic impairment^7,9–12^. Thus, identifying bioactive signals capable of modifying this hypothalamic neuroimmune environment may reveal previously unrecognized modulators of neuroendocrine energy regulation.

One potential source of such signals is platelet-rich plasma (PRP), an autologous blood-derived preparation enriched in platelets and plasma components. Platelets can release a broad repertoire of bioactive molecules, including platelet-derived growth factor, transforming growth factor-β, vascular endothelial growth factor, insulin-like growth factor-1, cytokines, and chemokines. Collectively, these signals regulate cell proliferation, angiogenesis, extracellular matrix remodeling, and immune responses^13–15^. Their abundance and biological availability vary with platelet concentration, leukocyte content, preparation method, and activation status. PRP is therefore best considered a heterogeneous source of interacting biological signals rather than a uniformly anti-inflammatory agent with a single mechanism of action. Given the ability of its components to regulate cellular and immune processes, platelet-derived signals could plausibly influence glial and neural states when present in the central nervous system. Whether the hypothalamic neuroendocrine system responds to this complex signaling milieu, however, is unknown.

Support for neural responsiveness to PRP comes primarily from models of neural injury. PRP improved locomotor recovery and promoted angiogenesis and tissue repair in experimental spinal cord injury^16,17^, promoted axonal growth in a brain-spinal cord organ coculture system^18^, and has been investigated in models of cerebral ischemia, traumatic brain injury, and peripheral nerve injury^19–21^. However, these studies largely focus on regeneration after injury rather than hypothalamic neuroendocrine function or homeostatic control of metabolism. It therefore remains unclear whether PRP can influence hypothalamic cellular responses and systemic energy balance in the absence of direct neural injury.

To address this gap, we used central administration of autologous PRP as a proof-of-concept approach to test whether the brain is responsive to platelet-derived signals in two complementary settings. Lipopolysaccharide (LPS)-induced systemic inflammation was used to examine hypothalamic microglial responses during an acute inflammatory challenge, whereas diet-induced obesity (DIO) was used to examine responses during chronic metabolic stress and established obesity. We hypothesized that central PRP administration would modify hypothalamic microglial status and alter whole-body energy balance. To test this hypothesis, we combined region-specific microglial analysis with comprehensive metabolic phenotyping, whole-hypothalamus RNA sequencing, and qPCR analyses to define the cellular, physiological, and molecular responses to central PRP administration.

## Materials and Methods

### 2.1. Animals

Male C57BL/6J mice (#000664, Jackson Laboratory, Bar Harbor, ME, USA) were used in this study. All animal procedures were reviewed and approved by the Institutional Animal Care and Use Committee of the University of Illinois Chicago. This proof-of-concept study was conducted in male mice. Whether the findings generalize to females remains to be determined. Mice were housed in a temperature- and humidity-controlled environment (23 ± 3°C and 70 ± 10% humidity) under a 14:10 h light/dark cycle, with ad libitum access to food and water. Mice were fed either a standard chow diet (CD; Teklad LM-485 Mouse/Rat Sterilizable Diet, #7012, Envigo, Indianapolis, IN, USA; 17% kcal from fat, 25% kcal from protein, and 58% kcal from carbohydrate) or a high-fat diet (HFD; Research Diets D12492, Rodent Diet With 60 kcal% Fat, Research Diets, Inc., New Brunswick, NJ, USA).

### 2.2. Preparation of PRP and Artificial Cerebrospinal Fluid

Autologous PRP was prepared from approximately 300 μL of whole blood collected from the facial vein of each mouse. For the DIO experiment, blood was collected at week 8 of HFD feeding, approximately 4 weeks before ICV administration. For the LPS experiment, blood was also collected approximately 4 weeks before ICV administration. aCSF-treated control mice underwent the same blood collection, restraint, and recovery procedures as PRP-treated mice. Whole blood was collected into tubes pretreated with 0.5 M EDTA, pH 8.0 (Research Products International, Mt. Prospect, IL, USA; Cat. No. E14000-100.0), as an anticoagulant and centrifuged at 300 × g for 10 min at 4°C. Approximately 150 μL of the upper platelet-containing plasma fraction was carefully collected and centrifuged at 800 × g for 15 min at 4°C to pellet the platelets.

After removal of the supernatant, the platelet pellet was gently resuspended in aCSF at a volume corresponding to approximately one-eighth of the initial whole-blood volume to generate concentrated PRP. The resulting autologous PRP was stored at −80°C until administration. aCSF was used for PRP resuspension in the experimental cohorts so that PRP-treated and control mice received the same intracerebroventricular vehicle and to minimize perturbation of the cerebrospinal fluid ionic environment. Artificial cerebrospinal fluid (aCSF) was prepared in sterile distilled water and contained 148 mM NaCl, 3.0 mM KCl, 1.4 mM CaCl₂, 0.8 mM MgCl₂, 0.8 mM Na₂HPO₄, and 0.2 mM NaH₂PO₄. The solution was sterilized by passage through a 0.22-μm membrane filter and stored at 4°C until use.

To characterize the platelet concentration achieved by this preparation procedure, PRP was subsequently prepared from an independent cohort of six mice using the same centrifugation protocol. For this validation cohort, platelet pellets were resuspended in calcium- and magnesium-free 1× PBS at the same one-eighth-volume ratio to facilitate hematology-based platelet counting. Platelet concentrations were measured directly using an ADVIA 120 automated hematology analyzer (Siemens Healthcare Diagnostics Inc., Tarrytown, NY, USA) based on two-angle laser light-scatter optical analysis. Because platelet recovery during collection of platelet-containing plasma varied among samples, the actual platelet concentration was not expected to correspond uniformly to the theoretical eightfold volumetric concentration. In this validation cohort, the measured PRP platelet concentration was 4.30 ± 1.08 × 10^6^ platelets/μL (mean ± SEM, n = 6), with individual values ranging from 2.11 to 9.31 × 10^6^ platelets/μL. These measurements were used to estimate the platelet concentration of the PRP administered in the original experiments, corresponding to an estimated mean dose of approximately 4.30 × 10^6^ platelets per 1-μL injection.

### 2.3. Intracerebroventricular Administration

PRP was administered into the right lateral ventricle using either an indwelling ICV cannula or direct stereotaxic injection, depending on the experimental purpose. aCSF was administered using the corresponding procedure as the vehicle control.

#### 2.3.1. Implanted-cannula administration

This procedure was used for DIO Cohort 1. ICV administration was performed through an indwelling guide cannula. Mice underwent stereotaxic surgery under isoflurane anesthesia using an Angle Two Small Animal Stereotaxic Instrument (Leica Biosystems, Nussloch, Baden-Württemberg, Germany). An indwelling 26-gauge ICV guide cannula (OD 0.48 mm; Cat. No. 62003, RWD Life Science, Sugar Land, TX, USA) with a 2.3-mm projection was unilaterally implanted into the right lateral ventricle at the following coordinates relative to bregma: AP, −0.35 mm; ML, +1.0 mm; and DV, −2.5 mm. The cannula was anchored to the skull using dental cement. Correct cannula placement and patency were assessed 3–4 days after surgery by the induction of a rapid drinking response following ICV administration of angiotensin II (10 ng), as previously described ^22^. Mice with misplaced or blocked cannulas were excluded from subsequent analyses. The remaining mice were allowed to recover, and PRP or aCSF was administered approximately 2 weeks after cannulation. For ICV administration, an internal injector (OD 0.30 mm; Cat. No. 62203, RWD Life Science, Sugar Land, TX, USA) extending 0.5 mm beyond the tip of the guide cannula was used.

#### 2.3.2. Direct stereotaxic ICV injection

This procedure was used for the LPS cohort and DIO Cohort 2. PRP or aCSF was administered by direct stereotaxic injection under isoflurane anesthesia using the same stereotaxic apparatus. The injector was positioned in the right lateral ventricle at the following coordinates relative to bregma: AP, −0.35 mm; ML, +1.0 mm; and DV, −2.5 mm. A total volume of 1 μL was delivered at a rate of 0.5 μL/min using a 2-μL Neuros Syringe, Model 7002 (Cat. No. 65459-01; Hamilton Company, Reno, NV, USA). The injector was left in place for 2 min after injection before being slowly withdrawn.

### 2.4 Diet-Induced Obesity and PRP ICV Administration

Male mice were placed on an HFD beginning at 8 weeks of age, with body weight and food intake monitored weekly. Two independent cohorts were used to assess longitudinal metabolic outcomes and short-term hypothalamic molecular responses, respectively. For both cohorts, approximately 300 μL of whole blood was collected from each mouse at week 8 of HFD feeding, approximately 4 weeks before PRP or aCSF administration. aCSF-treated control mice underwent the same blood collection, restraint, and recovery procedures as PRP-treated mice.

#### Cohort 1: Longitudinal metabolic phenotyping

Mice underwent ICV cannulation after 10 weeks of HFD feeding, at approximately 18 weeks of age. At the time of cannulation, mice weighed 34.1 ± 1.6 g (mean ± SEM). Following cannulation, mice were allowed to recover for approximately 2 weeks before treatment. At week 12 of HFD feeding, mice received a single 1-μL ICV injection of PRP or aCSF through the indwelling guide cannula (n = 7 per treatment group), approximately 4 h after lights on (approximately 10:00 a.m.). Thus, PRP/aCSF administration occurred approximately 4 weeks after blood collection and 2 weeks after cannulation.

Metabolic cage analysis was performed on post-injection day 7, followed by GTT on day 14, and ITT on day 21. On post-injection day 28, mice were fasted overnight, and fasting blood glucose was measured immediately before terminal tissue collection at approximately 4 h after lights on (approximately 10:00 a.m.). Mice were deeply anesthetized with inhaled isoflurane and transcardially perfused with PBS followed by 10% neutral buffered formalin for brain collection. Brains were post-fixed overnight at 4°C in a 1:1 mixture of 10% formalin and 20% sucrose, cryoprotected in 20% sucrose (w/v) at 4°C, and sectioned at 30 μm for subsequent immunofluorescence staining. Gonadal white adipose tissue (gWAT), inguinal white adipose tissue (iWAT), interscapular brown adipose tissue (BAT), liver, testes, gastrocnemius, and tibialis anterior muscles were also collected.

#### Cohort 2: Short-term histological and molecular analyses

An independent cohort of DIO mice was used to examine short-term hypothalamic responses to PRP. Mice were approximately 20 weeks of age and weighed 34.2 ± 1.3 g (mean ± SEM) at the time of treatment. After 12 weeks of HFD feeding, mice received a direct stereotaxic ICV injection of 1 μL PRP or aCSF (n = 10 per treatment group) into the right lateral ventricle without prior cannula implantation. Body weight and food intake were recorded daily for 7 days after injection.

On post-injection day 7, mice were fasted for overnight before tissue collection at approximately 4 h after lights on (approximately 10:00 a.m.). Within each treatment group, six mice were used for molecular analyses, and four mice were used for histological analyses. For molecular analyses, mice were euthanized by carbon dioxide asphyxiation followed by cervical dislocation, and hypothalamic tissues were rapidly dissected, frozen on dry ice, and stored at −80°C until quantitative PCR or RNA sequencing analysis. For histological analyses, mice were deeply anesthetized with inhaled isoflurane and transcardially perfused with PBS followed by 10% neutral buffered formalin. Brains were collected, post-fixed overnight at 4°C in a 1:1 mixture of 10% formalin and 20% sucrose, cryoprotected in 20% sucrose (w/v) at 4°C, and sectioned at 30 μm for immunofluorescence staining.

An indwelling guide cannula was used in Cohort 1 to allow full recovery from stereotaxic surgery before PRP administration and subsequent longitudinal metabolic phenotyping, thereby minimizing the influence of acute surgical stress on metabolic outcomes. In contrast, direct stereotaxic injection was used in Cohort 2 because this cohort was designed for short-term hypothalamic molecular and histological analyses, and avoiding chronic cannula implantation minimized potential local tissue perturbation that could confound these readouts.

### 2.5 LPS-induced Systemic Inflammation and PRP ICV Injection

Male mice approximately 12 weeks of age and weighing 23.6 ± 0.6 g (mean ± SEM) were allocated to three experimental groups based on baseline body weight to ensure comparable body weights among groups before treatment: Saline+aCSF, LPS+aCSF, and LPS+PRP (n = 4/4/4). Baseline body weight did not differ significantly among the three groups. Mice received an intraperitoneal (i.p.) injection of LPS from Escherichia coli O55:B5 (#HY-D1056, MedChemExpress, Monmouth Junction, NJ, USA), dissolved in sterile saline at a concentration of 0.2 mg/mL and administered at 1 mg/kg body weight in an injection volume of 5 μL/g, or an equivalent volume of sterile saline. LPS or saline was administered at approximately 4 h after light on (approximately 10:00 a.m.). 24 hours later, body weight was recorded, followed by ICV injection of PRP or aCSF as described in Section 2.3. Twenty-four hours after ICV injection, body weight was measured again. Mice were maintained with ad libitum access to food and water until terminal tissue collection, which was performed at approximately 10:00 a.m. Mice were then deeply anesthetized and transcardially perfused, and brains were processed for immunofluorescence analysis as described in Sections 2.8.

### 2.6 Metabolic Phenotyping

Whole-body energy metabolism was assessed with a Promethion metabolic phenotyping system (Sable Systems International, Las Vegas, NV, USA) in the Metabolic Phenotyping Core at the Biologic Resources Laboratory, University of Illinois Chicago. Animals were housed individually in metabolic chambers with unrestricted access to food and water and were allowed to acclimate to the system for 72 h before measurements were analyzed. Chamber temperature, humidity, and the light–dark schedule were maintained under standardized conditions throughout the experiment. The system was calibrated and operated in accordance with the manufacturer’s recommendations and established core procedures.

Raw calorimetry data were analyzed using ExpeData software (Sable Systems) together with CalR (https://calrapp.org/). Fat and carbohydrate utilization were calculated from oxygen consumption (VO₂) and carbon dioxide production (VCO₂) using the following equations: fat utilization = 1.67 × VO₂ − 1.67 × VCO₂, and carbohydrate utilization = 4.55 × VCO₂ − 3.21 × VO₂.

Differences in energy expenditure between groups were evaluated by ANCOVA using the MMPC Energy Expenditure Analysis platform (http://www.mmpc.org/shared/regression.aspx), with total daily energy expenditure modeled as the outcome variable and body weight included as a covariate. Treatment-by-body weight interactions were tested to assess the homogeneity of regression slopes. When the interaction was not significant, a common-slope ANCOVA model was used to estimate the body weight– adjusted treatment effect. When a significant treatment-by-body weight interaction was detected, indicating that the relationship between energy expenditure and body weight differed between treatment groups, a single common adjusted treatment effect was not reported. Instead, the interaction was interpreted on the basis of the treatment-specific regression slopes and fitted regression lines. A secondary ANCOVA model included physical activity as an additional covariate to assess whether the treatment-associated difference in energy expenditure was explained by locomotor activity. Additional details of the statistical models are specified in the relevant figure legends.

### 2.7 Glucose homeostasis tests

For the glucose tolerance test (GTT), mice were fasted overnight for 13 h (19:00–08:00), with free access to water. Blood glucose was measured from tail-vein blood using a StatStrip Xpress 2 Glucose Meter (Cat. No. 56510, Nova Biomedical, Waltham, MA, USA). After determination of baseline glucose levels, mice were administered dextrose (D9434, Sigma-Aldrich, St. Louis, MO, USA) by i.p. injection at 2 g/kg body weight. Dextrose was prepared at a concentration of 0.4 g/mL and administered at an injection volume of 5 μL/g body weight. Blood glucose concentrations were subsequently measured at 15, 30, 60, and 120 min after injection.

For the insulin tolerance test (ITT), mice were fasted for 4 h beginning at 09:00, with water available throughout the fasting period. Following baseline glucose measurement, insulin (Cat. No. 0002-75-1001, Eli Lilly and Company, Indianapolis, IN, USA) was prepared at a concentration of 1.4 U/mL and administered by intraperitoneal injection at 0.7 U/kg body weight, corresponding to an injection volume of 0.5 μL/g body weight. Blood glucose levels were then measured at 15, 30, 60, and 120 min after insulin administration. Integrated glucose responses during GTT and ITT were summarized by area under the curve and compared between treatment groups.

### 2.8 Immunofluorescence Staining

Fixed brains were sectioned coronally at a thickness of 30 μm using an HM 430 Sliding Microtome (Epredia). Free-floating sections were blocked and permeabilized for 1 h at room temperature in PBS containing 5% normal goat serum and 0.3% Triton X-100, followed by overnight incubation at 4°C with rabbit anti-IBA1 antibody (1:1,000; Fujifilm Wako Pure Chemical Corporation, Osaka, Japan; Cat. No. 019-19741). After washing, sections were incubated with Alexa Fluor 488-conjugated goat anti-rabbit secondary antibody (1:500; Jackson Immuno Research, West Grove, PA, USA; Cat. No. 111-545-003) for 2 h at room temperature and counterstained with DAPI.

Fluorescence images were acquired using an Olympus BX51 fluorescence microscope. Images were analyzed using ImageJ version 1.54p. Images were converted to 8-bit, followed by background subtraction and thresholding. IBA1-positive cell number and mean particle area were quantified using the Analyze Particles function in ImageJ ^23^.

Quantification was performed within predefined regions of interest in the arcuate nucleus (ARC), ventromedial hypothalamus (VMH), and dorsomedial hypothalamus (DMH). Sections spanning approximately −1.70 to −2.30 mm from bregma were analyzed. For each mouse, five sections per brain region were selected at approximately 150-μm intervals. Measurements from the five sections within each region were averaged to obtain a single value per mouse. The mouse was the biological experimental unit..

### 2.9 RNA Sequencing and Bioinformatic Analysis

Purified total RNA from hypothalamic tissues was submitted to Plasmidsaurus (Eugene, OR, USA) for RNA sequencing. Sequencing was performed as stranded single-end 3′-end RNA sequencing with a read length of approximately 90 bp. According to the Plasmidsaurus RNA-seq workflow, the service is designed to yield approximately 20 million raw reads and approximately 10 million UMI-deduplicated reads per sample. In the present study, successful sequencing runs generated approximately 18.7–20.6 million raw reads per run. Following processing and merging of sequencing runs where applicable, the final datasets contained approximately 21.7–24.0 million alignments per biological sample, with overall mapping rates of 95.5–98.0%. Gene-level quantification yielded approximately 12.1–14.1 million assigned reads per sample. Sequencing data were processed using the standard Plasmidsaurus RNA-seq pipeline. Reads were quality-filtered using FastP, aligned to the mouse reference genome using STAR, deduplicated based on UMIs using UMICollapse, and quantified at the gene level using featureCounts.

A total of 12 biological samples were included in the study. One sample with insufficient sequencing depth in the initial run was resequenced, and six additional samples were subjected to repeat sequencing to ensure adequate sequencing quality and technical reproducibility. For resequenced samples, data from the initial and repeat sequencing runs were merged at the sample level prior to downstream analysis. To account for technical variation across sequencing runs, batch effects were corrected using the APlasmidsaurus anchor-based batch-correction workflow implemented with RUVseq, with technical replicates across sequencing runs serving as anchor samples. Resequencing did not constitute additional biological replicates.

Differential gene expression between the PRP- and aCSF-treated groups was analyzed using the Plasmidsaurus bioinformatics pipeline, and differential expression results were visualized using volcano plots. Gene set enrichment analysis (GSEA) was performed using the MSigDB Hallmark gene sets and KEGG pathway gene sets to identify biological pathways differentially enriched between treatment groups. Enrichment was summarized using normalized enrichment scores (NES), with positive and negative NES values indicating pathways enriched in the PRP- and aCSF-treated groups, respectively.

### 2.10 RNA Isolation and Quantitative PCR

Total RNA was isolated from hypothalamic tissues from DIO Cohort 2 (n = 6 per treatment group) using TRIzol reagent (Invitrogen, Thermo Fisher Scientific, Waltham, MA, USA; Cat. No. 15596018) according to the manufacturer’s instructions. RNA was reverse-transcribed into cDNA using the High-Capacity cDNA Reverse Transcription Kit with RNase Inhibitor (Applied Biosystems, Thermo Fisher Scientific, Waltham, MA, USA; Cat. No. 4374967). qPCR was performed using 2X Universal SYBR Green Fast qPCR Mix (ABclonal, Woburn, MA, USA; Cat. No. RK21203) according to the manufacturer’s instructions. Relative gene expression was normalized to *Gapdh* and calculated using the 2^−ΔΔCt method with the aCSF group as calibrator. Primer sequences are provided in Table S1. The qPCR assays used samples from the same biological cohort as RNA sequencing and therefore supported selected RNA-seq-associated changes rather than constituting an independent biological replication.

### 2.11 Statistical Analysis

Statistical analyses were performed using GraphPad Prism 10 (GraphPad Software, San Diego, CA, USA), except for the calorimetry ANCOVA and RNA-seq analyses described above. The mouse was the biological experimental unit. Data normality was assessed before parametric testing. Comparisons between two groups were analyzed using two-tailed unpaired Student’s *t*-test, whereas dynamic changes over time or differences among multiple groups were evaluated using one-way or two-way analysis of variance (ANOVA) followed by Tukey’s or Sidak’s *post hoc* test. All data are presented as mean ± SEM, and a *P* value < 0.05 was considered statistically significant.

## Results

### 3.1 Central PRP altered metabolic phenotypes in diet-induced obese mice

To investigate the metabolic effects of central PRP administration in diet-induced obesity, male mice were fed an HFD beginning at 8 weeks of age. After 10 weeks of HFD feeding, mice underwent ICV cannulation and were individually housed. PRP or aCSF was administered through the implanted guide cannula after 12 weeks of HFD feeding. Metabolic cage analysis, GTT, and ITT were subsequently performed, followed by tissue collection after 16 weeks of HFD feeding (Figure 1A).

**Figure 1.**
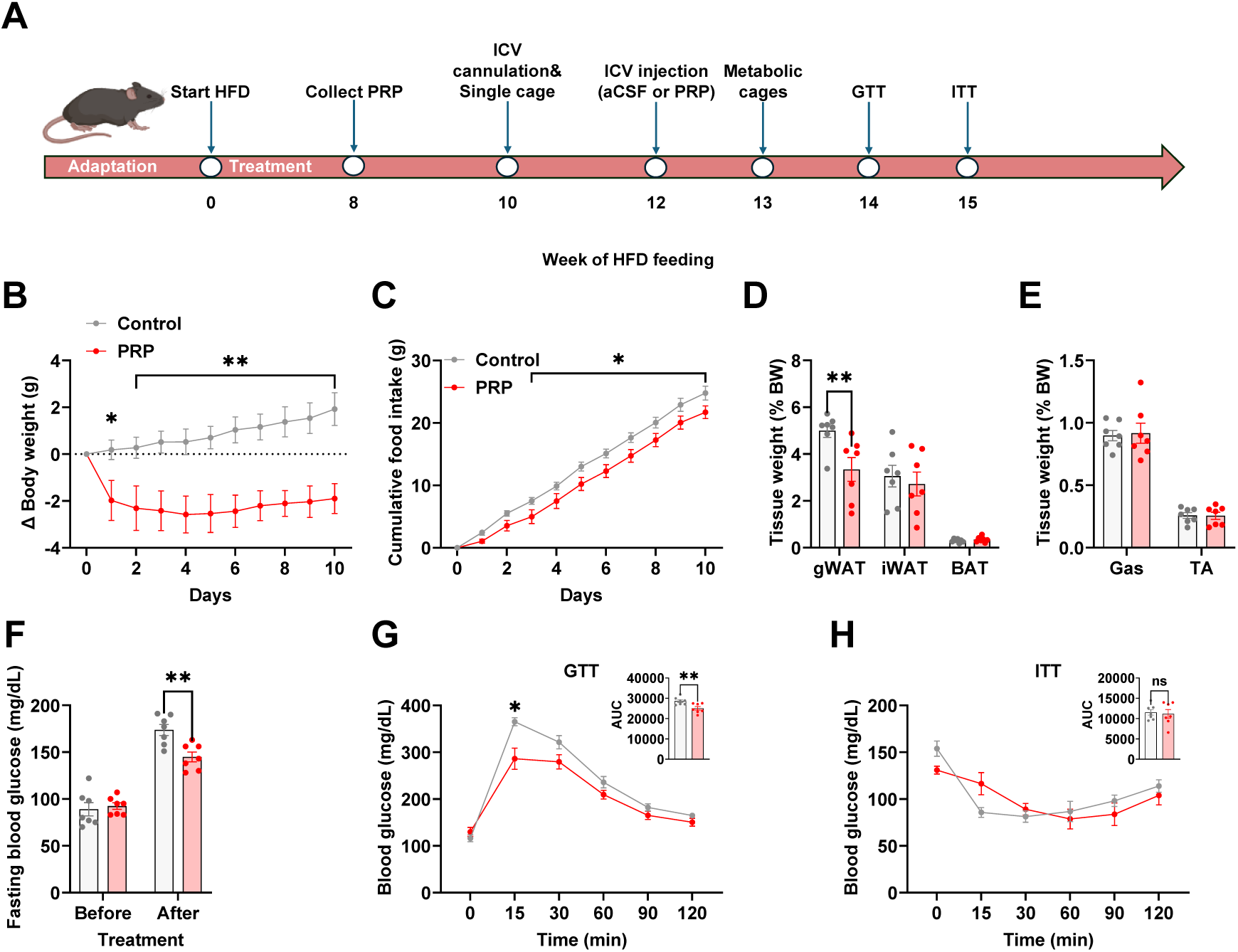
Effects of central PRP on metabolic phenotypes in diet-induced obese mice. **(A)** Experimental schematic showing the initiation of HFD feeding, autologous PRP collection, ICV cannulation, ICV administration of aCSF or PRP, metabolic cage analysis, glucose tolerance test (GTT), and insulin tolerance test (ITT). **(B)** Body weight changes over 10 days after ICV administration. **(C)** Cumulative food intake over 10 days after ICV administration. **(D)** Relative weights (tissue weight normalized to body weight) of gonadal white adipose tissue (gWAT), inguinal white adipose tissue (iWAT), and brown adipose tissue (BAT). **(E)** Relative weights of gastrocnemius (Gas) and tibialis anterior (TA) muscles. **(F)** Fasting blood glucose levels before and after ICV administration. **(G)** Blood glucose concentrations during the GTT and corresponding area under the curve (AUC). **(H)** Blood glucose concentrations during the ITT and corresponding AUC. Data are presented as mean ± SEM. n values indicate biological replicates (individual mice): n = 7 mice per group. Statistical significance was analyzed by two-way ANOVA with Sidak’s multiple-comparisons test (B–H), and AUC values in (G, H) were compared using unpaired two-tailed Student’s *t*-tests. Significance thresholds: \**P* < 0.05, \*\**P* < 0.01.

Following ICV administration, body weight gradually increased in control mice, whereas it continued to decrease in PRP-treated mice, although the rate of decline became slower over time. Body weight change remained lower in the PRP group throughout the monitoring period (Figure 1B). Cumulative food intake was also lower in PRP-treated mice, with significant differences observed from day 3 through day 10 (Figure 1C).

The relative weight of gonadal white adipose tissue (gWAT) was significantly lower in the PRP group than in the control group. No significant differences were observed in inguinal white adipose tissue (iWAT) or brown adipose tissue (BAT) between the two groups (Figure 1D). Similarly, gastrocnemius and tibialis anterior muscle weights were not significantly different between the groups (Figure 1E).

Before treatment, fasting blood glucose levels were comparable between the two groups. After treatment, fasting blood glucose was significantly lower in PRP-treated mice than in control mice (Figure 1F). During the glucose tolerance test (GTT), blood glucose was significantly lower in the PRP group at 15 min after glucose administration, with no significant differences at the remaining time points. The GTT AUC was also significantly lower in the PRP group (Figure 1G). During the insulin tolerance test (ITT), blood glucose levels did not differ significantly between the control and PRP groups at any time point, and the ITT AUC was also not significantly different between the groups (Figure 1H).

In a separate cohort used for short-term analyses, PRP was administered by direct ICV injection without cannulation. The PRP group showed a greater reduction in body weight than the control group throughout the 7-day observation period. Cumulative food intake was slightly lower in the PRP group, with a significant difference observed on day 7 (Figure S1).

### 3.2 Central PRP increased energy expenditure and physical activity in diet-induced obese mice

To test whether central PRP affects energy metabolism, we access energy metabolism through Sable Promethion system. Energy expenditure was generally higher in PRP-treated mice over the 24-h recording period (Figure 2A). Phase-based analysis showed significantly higher energy expenditure in the PRP group during both the light and dark phases, as well as across the entire 24-h period (Figure 2B).

**Figure 2.**
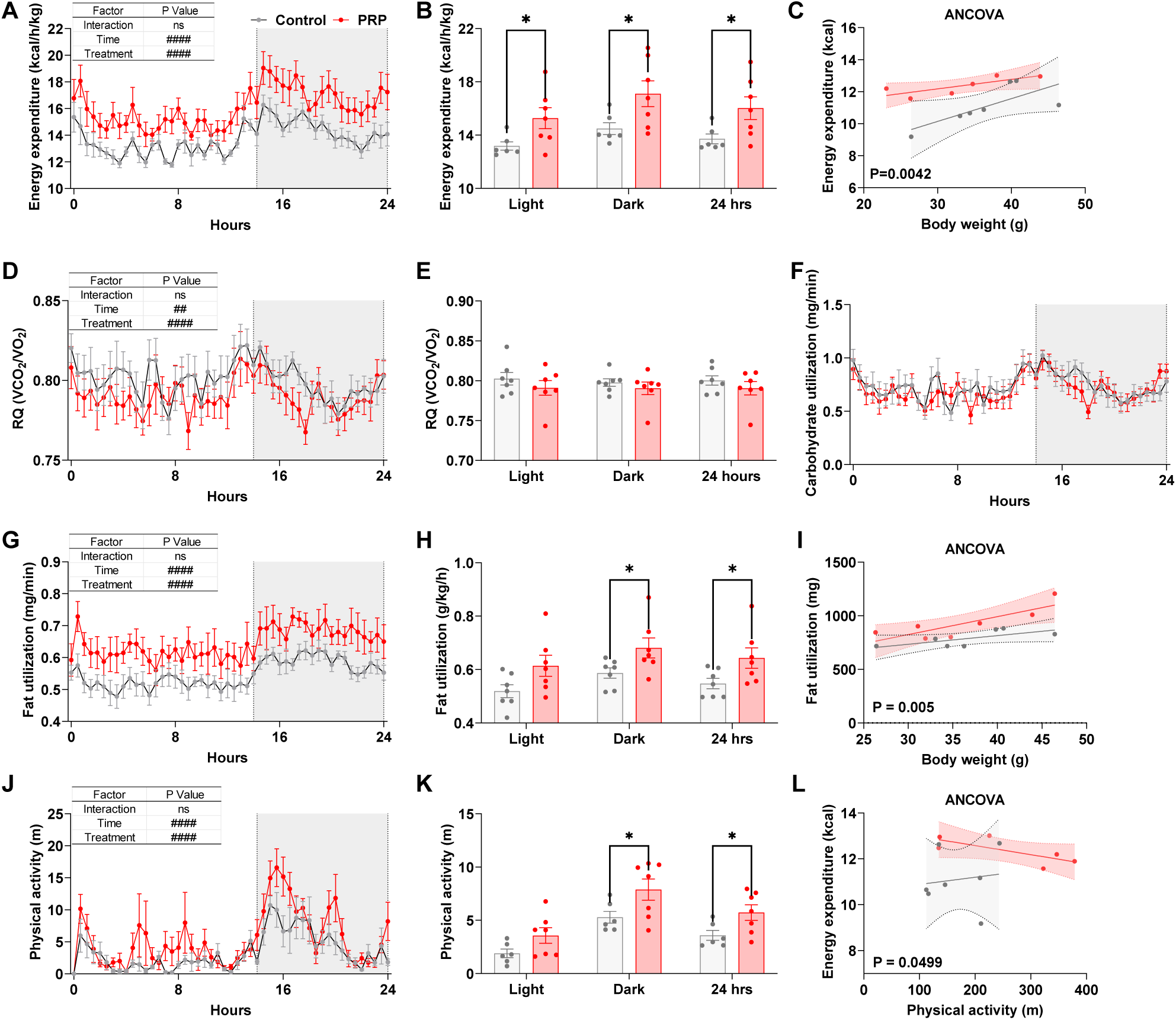
Effects of central PRP on whole-body energy metabolism in diet-induced obese mice. **(A)** Energy expenditure over the 24-h recording period. **(B)** Average energy expenditure during the light phase, dark phase, and entire 24-h period. **(C)** ANCOVA of energy expenditure with body weight as a covariate. **(D)** Respiratory quotient (RQ) over the 24-h recording period. **(E)** Average RQ during the light phase, dark phase, and entire 24-h period. **(F)** Carbohydrate utilization over the 24-h recording period. **(G)** Fat utilization over the 24-h recording period. **(H)** Average fat utilization during the light phase, dark phase, and entire 24-h period. **(I)** ANCOVA of fat utilization with body weight as a covariate. **(J)** Physical activity over the 24-h recording period. **(K)** Average physical activity during the light phase, dark phase, and entire 24-h period. **(L)** Relationship between physical activity and energy expenditure analyzed by ANCOVA. Data are presented as mean ± SEM; n = 7 mice per group. Statistical significance was analyzed by two-way ANOVA with Sidak’s multiple-comparisons test (A, B, D, G, H, J, K). Energy expenditure in (C) or fat utilization (I) was analyzed by ANCOVA with body weight as a covariate; the treatment × body weight interaction was not significant, whereas the treatment effect remained significant after adjustment for body weight (*P* < 0.05). The relationship between physical activity and energy expenditure in (L) was analyzed by ANCOVA. Significance thresholds: ##*P* < 0.05, ####*P* < 0.0001, \**P < 0.05*; ns = not significant.

ANCOVA revealed no significant treatment-by-body weight interaction, indicating that the relationship between body weight and energy expenditure did not significantly differ between the control and PRP groups. After accounting for body weight, energy expenditure differed significantly between the two groups (treatment effect, *P* =0.0042; Figure 2C).

Respiratory quotient (RQ) was modest but consistently lower in PRP-treated mice across the 24-h recording period (Figure 2D). Two-way ANOVA revealed significant main effects of time and treatment, with no significant time-by-treatment interaction. When RQ was averaged by light phase, dark phase, or over the entire 24-h period, no significant differences were detected between groups (Figure 2E).

We next examined substrate utilization to determine whether PRP treatment altered metabolic fuel preference. Carbohydrate utilization did not differ significantly between control and PRP-treated mice across the 24-h recording period (Figure 2F). In contrast, fat utilization was significantly increased in PRP-treated mice across the 24-h period (Figure 2G). Two-way ANOVA revealed significant main effects of time and treatment, with no significant time-by-treatment interaction. Phase-based analysis showed significantly greater fat utilization in PRP-treated mice during the dark phase and across the entire 24-h period, whereas the difference during the light phase was not significant (Figure 2H). ANCOVA further showed that the increase in fat utilization remained significant after adjustment for body weight (P = 0.005; Figure 2I).

Finally, we examined whether PRP treatment altered physical activity and whether increased activity could account for the elevation in energy expenditure. Physical activity was significantly higher in PRP-treated mice across the 24-h recording period (Figure 2J and 2K). ANCOVA revealed no significant treatment-by-physical activity interaction, indicating similar relationships between physical activity and energy expenditure between groups. Importantly, energy expenditure remained significantly higher in PRP-treated mice after adjustment for physical activity (Figure 2L), suggesting that the increase in physical activity does not fully account for the PRP-associated elevation in energy expenditure.

### 3.3 Central PRP reduced microglial cell size in the ARC of diet-induced obese mice

To determine whether central PRP administration altered hypothalamic microglia in diet-induced obese mice, IBA1 immunofluorescence staining was performed in the ARC, VMH, and DMH. The number of IBA1-positive cells was not significantly different between the control and PRP groups in the ARC, VMH, or DMH (Figure 3A and 3B). In contrast, IBA1-positive cell size was significantly reduced in the ARC of PRP-treated mice compared with control mice. IBA1-positive cell size also appeared lower in the VMH and DMH, although these differences were not statistically significant (Figure 3C).

**Figure 3.**
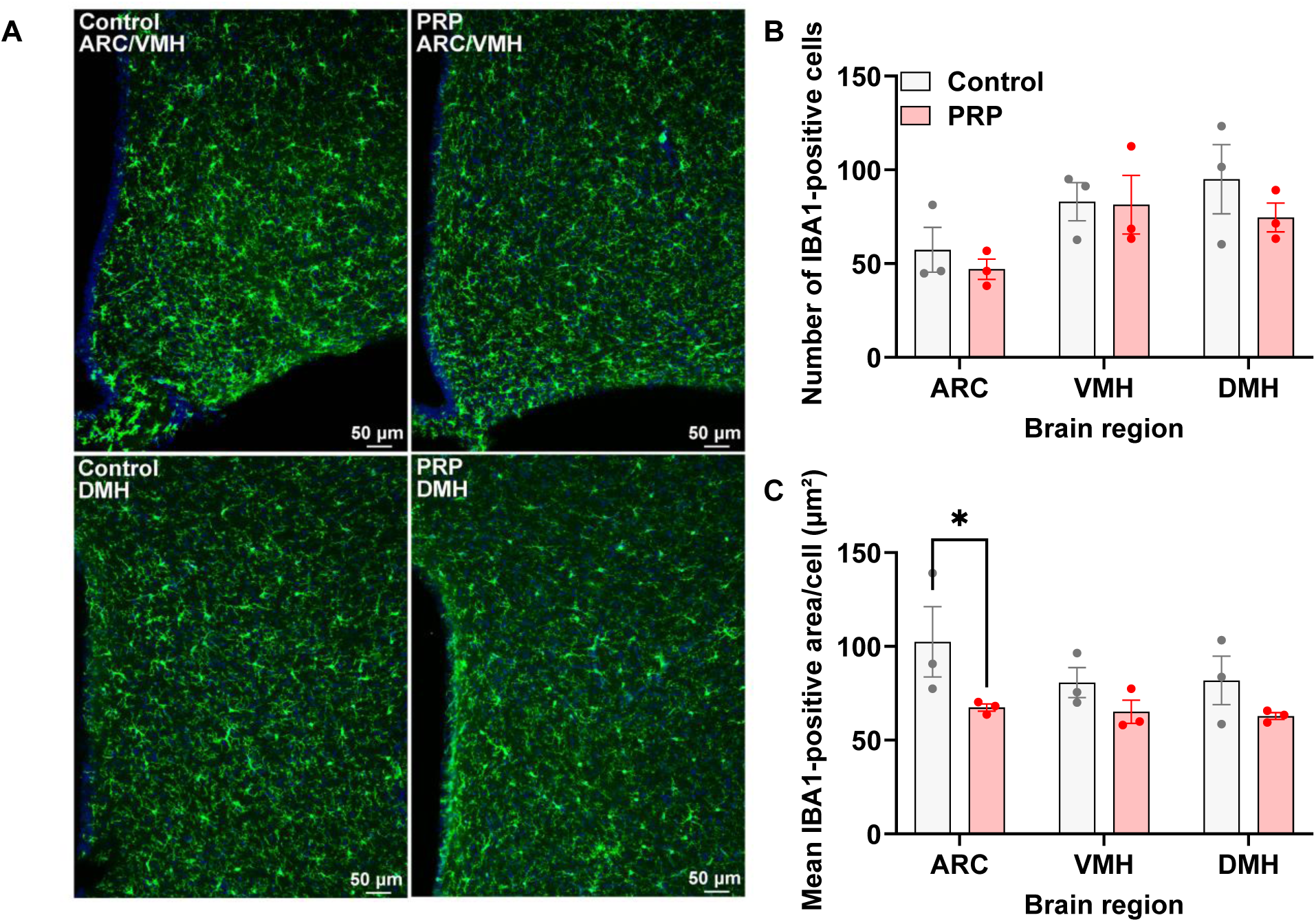
Effects of central PRP on hypothalamic microglial morphology in diet-induced obese mice. **(A)** Representative IBA1 (green) immunofluorescence images of the ARC/VMH and DMH in control and PRP-treated mice. Nuclei were labeled with DAPI (blue). **(B)** IBA1-positive cell number in the ARC, VMH, and DMH. **(C)** Mean IBA1-positive area per cell in the ARC, VMH, and DMH. Data are presented as mean ± SEM; n = 3 mice per group. Statistical significance was analyzed by two-way ANOVA with Sidak’s multiple-comparisons test (B, C). Significance thresholds: \**P* < 0.05.

These results indicate that central PRP administration altered microglial morphology in a region-specific manner, with a significant reduction in IBA1-positive cell size in the ARC but no detectable change in IBA1-positive cell number.

### 3.4 Central PRP partially attenuated LPS-induced changes in hypothalamic microglial morphology

To determine whether the hypothalamic microglial morphological changes observed in DIO mice were also present under acute inflammatory conditions, we examined the effects of central PRP administration in an LPS-induced systemic inflammation model. Mice received an intraperitoneal injection of saline or LPS on day 1, followed by an ICV injection of aCSF or PRP on day 2. Brain tissues were collected on day 3 for IBA1 immunofluorescence staining to access the number and morphology of microglia (Figure 4A).

**Figure 4.**
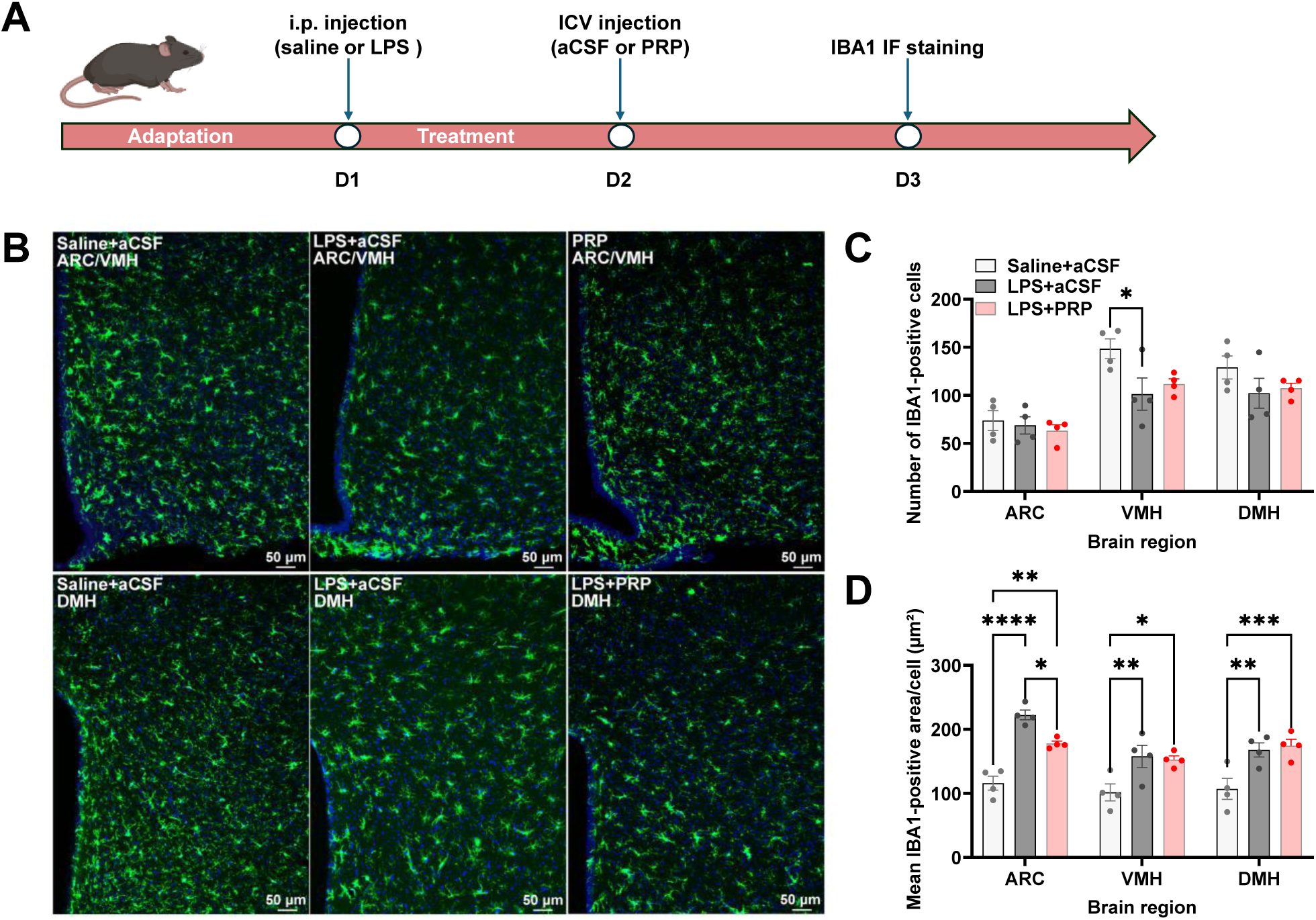
Effects of central PRP on hypothalamic microglial morphology in LPS-treated mice. **(A)** Experimental timeline showing intraperitoneal administration of saline or LPS, followed by ICV administration of aCSF or PRP and subsequent tissue collection for IBA1 immunofluorescence staining. **(B)** Representative IBA1 immunofluorescence images of the ARC/VMH and DMH in Saline+aCSF, LPS+aCSF, and LPS+PRP groups. **(C)** IBA1-positive cell number in the ARC, VMH, and DMH. **(D)** Mean IBA1-positive area per cell in the ARC, VMH, and DMH. Data are presented as mean ± SEM; n = 4 mice per group. Statistical significance was analyzed by two-way ANOVA with Sidak’s multiple-comparisons test (C, D). Significance thresholds: \**P* < 0.05, \*\**P* < 0.01, \*\*\**P* < 0.001, \*\*\*\**P* < 0.0001.

PRP did not significantly alter LPS-induced body weight changes, as body weight loss was comparable between the LPS+aCSF and LPS+PRP groups. However, mice in the LPS+PRP group exhibited greater body weight loss than those in the Saline+aCSF group (Figure S3).

PRP did not significantly affect the number of IBA1-positive cells in the ARC, VMH, or DMH compared with the LPS+aCSF group. LPS treatment itself was associated with a reduction in IBA1-positive cell number in the VMH, whereas no significant group differences were observed in the ARC or DMH (Figure 4C). However, PRP selectively attenuated the LPS-induced increase in IBA1-positive cell size in the ARC, whereas no significant effect was observed in the VMH or DMH. Although PRP reduced cell size in the ARC compared with the LPS+aCSF group, values remained elevated relative to the Saline+aCSF group (Figure 4D).

These findings suggest that central PRP partially reduced the LPS-induced increase in microglial cell size in the ARC but did not significantly affect IBA1-positive cell number.

### 3.5 Central PRP altered hypothalamic gene expression and biological pathways in diet-induced obese mice

To investigate the molecular changes associated with central PRP administration, RNA sequencing was performed using hypothalamic tissues from control and PRP-treated mice. Differential expression analysis identified a larger number of upregulated genes than downregulated genes in the PRP group (Figure 5A).

**Figure 5.**
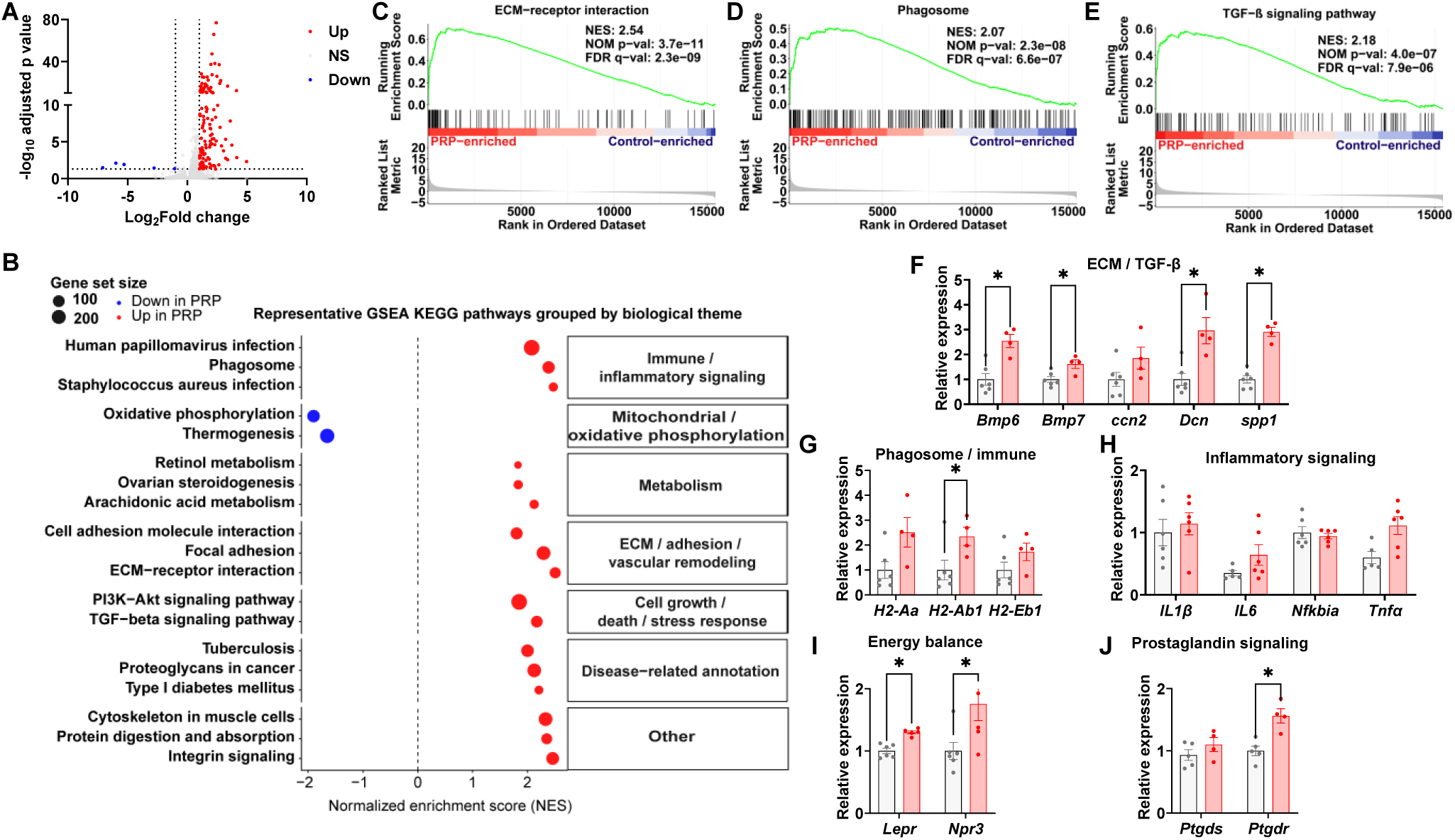
Effects of central PRP on hypothalamic gene expression and biological pathways in diet-induced obese mice. **(A)** Volcano plot of differentially expressed genes between control and PRP-treated mice. **(B)** Representative KEGG pathways identified by gene set enrichment analysis (GSEA) and grouped by biological theme. **(C–E)** Representative GSEA enrichment plots for ECM–receptor interaction (C), phagosome (D), and TGF-β signaling pathway (E). **(F)** Relative hypothalamic expression of genes associated with ECM/TGF-β-related signaling, including *Bmp6, Bmp7, Ccn2, Dcn,* and *Spp1*. **(G)** Relative expression of phagosome- and immune-related genes *H2-Aa, H2-Ab1,* and *H2-Eb1*. **(H)** Relative expression of inflammation-related genes *Il1β, Il6, Nfkbia,* and *Tnfα*. **(I)** Relative expression of the energy balance-related genes *Lepr* and *Npr3*. **(J)** Relative expression of the prostaglandin signaling-related genes Ptgds and Ptgdr. Data are presented as mean ± SEM; n = 6 mice per group. Statistical significance was assessed using unpaired two-tailed Student’s *t*-tests (F, G, I, J). Significance thresholds: \**P* < 0.05.

Gene set enrichment analysis (GSEA) of KEGG pathways revealed changes across several biological categories. Several pathways related to immune and inflammatory signaling, metabolism, extracellular matrix and cell adhesion, and cell growth and stress responses were positively enriched in the PRP group. In contrast, oxidative phosphorylation and thermogenesis pathways were negatively enriched (Figure 5B). Representative enrichment plots illustrated positive enrichment of the ECM-receptor interaction, phagosome, and TGF-β signaling pathways in the PRP group (Figure 5C–E).

Selected genes were further examined by qPCR. Among genes associated with ECM and TGF-β-related signaling, *Bmp6, Bmp7, Dcn,* and *Spp1* were significantly increased in the PRP group, whereas *Ccn2* was not significantly altered (Figure 5F). Among phagosome- and immune-related genes, *H2-Ab1* was significantly increased, whereas *H2-Aa* and *H2-Eb1* were unchanged (Figure 5G). Among inflammation-related genes, *Il1β, Il6, Nfkbia,* and *Tnfα* were not significantly altered (Figure 5H). The energy balance-related genes *Lepr* and *Npr3* were significantly increased in the PRP group (Figure 5I). Among prostaglandin signaling-related genes, *Ptgdr* was significantly increased, whereas *Ptgds* was unchanged (Figure 5J).

## Discussion

This study reveals a previously unrecognized capacity of centrally administered PRP to influence systemic energy homeostasis. Despite extensive investigation of PRP as a regenerative biologic, primarily in peripheral tissues and, within the central nervous system, in models of neural injury and repair^16–21^, its effects on hypothalamic control of metabolism remain largely undefined. Here, we show that a single intracerebroventricular PRP injection in mice with established DIO produced reproducible weight loss. This negative energy balance was accompanied by reduced food intake, increased energy expenditure and physical activity, decreased relative gWAT tissue weight, and lower fasting glucose. In parallel, PRP consistently reduced mean IBA1-positive area per cell in the ARC in both DIO and acute LPS-induced inflammation and elicited broad hypothalamic transcriptional changes. Together, these physiological, anatomical, and molecular findings identify central metabolic regulation as a previously underappreciated target of PRP.

The rationale for this conceptual framework is supported by evidence that centrally acting secreted signals can coordinate feeding, oxygen consumption, activity, and substrate utilization^24,25^, and that immune-related signals influence hypothalamic neurons, glia, and neurovascular cells^26–28^. Nevertheless, whether central exposure to PRP preparation can engage hypothalamic neuroendocrine system and influence whole-body metabolism has remained largely unexplored. The present study addresses this gap by linking central PRP exposure to an integrated metabolic phenotype and to cellular and molecular responses within the hypothalamus.

A key finding of this study is the magnitude and breadth of the response to a single central PRP injection. Body weight decreased and remained lower throughout the observation period, and this effect was replicated in an independent cohort, supporting the robustness and reproducibility of the response. In parallel, food intake decreased, whereas oxygen consumption, whole-body energy expenditure, fat utilization, and locomotor activity increased, indicating a coordinated shift in energy balance rather than an isolated change in any single endpoint. Importantly, energy expenditure remained elevated after adjustment for body weight. Because PRP also increased locomotor activity, we further assessed energy expenditure using physical activity as a covariate. The PA-by-treatment interaction was not significant, and the treatment effect on energy expenditure remained significant after adjustment for physical activity, although the effect was modest and close to the conventional significance threshold. These findings suggest that increased locomotor activity contributes to the elevated energy expenditure but does not fully account for it, raising the possibility that PRP also engages activity-independent components of energy expenditure. Further studies incorporating pair-feeding, direct assessment of thermogenesis, and experimental control of locomotor activity will be needed to define the relative contributions of these mechanisms.

Consistent with these whole-body phenotypes, the reductions in relative gWAT mass, fasting blood glucose, and integrated glucose excursion during the GTT suggest that central PRP signaling may drive peripheral metabolic benefits. Supporting this interpretation, iWAT, BAT, and skeletal muscle weights did not change significantly, implying that weight loss primarily reduced unhealthy abdominal fat while preserving subcutaneous/thermogenic depots and lean mass. Notably, glucose tolerance also improved, despite no detectable change in insulin sensitivity. This pattern is more consistent with improved glucose homeostasis, potentially via enhanced glucose-stimulated insulin secretion, than with increased insulin responsiveness. However, these data do not establish that gWAT loss was longitudinal, and the improved glycemic control could still be secondary to overall body weight loss. Future studies should therefore incorporate more precise longitudinal measurements of regional fat depots and assess glucose homeostasis before body weight divergence. Overall, the results support a model in which central reprogramming initiates weight and fat distribution changes that secondarily improve glycemic control, underscoring the need for baseline glucose characterization and longitudinal body-composition profiling.

A second major finding is that PRP elicited a convergent microglial response within the ARC across two markedly different biological contexts, acute systemic inflammation and chronic DIO. In both models, PRP reduced microglial soma area without altering microglial number, consistent with a shift in morphological state rather than changes in cellular abundance. The ARC is a key neuroendocrine interface that integrates hormonal and nutrient cues through circuits governing feeding and energy expenditure^1,2^, and the reproducibility of this microglial phenotype in the ARC therefore strengthens the biological significance of PRP’s central actions and nominates ARC as a candidate site for mechanistic interrogation. Although formal treatment-by-region interaction will be needed to establish regional selectivity, the ARC was the only region examined in which a statistically significant effect was detected, suggesting a degree of anatomical specificity. Notably, in obese mice the ARC microglial response coincided with reduced food intake and body weight, raising the possibility that PRP-responsive microglia influence neighboring metabolic neurons to shape energy balance, as demonstrated by many studies^29–32^. While the current data does not demonstrate causality between microglial remodeling and metabolic outcomes, the anatomical and physiological convergence supports a model in which PRP engages ARC microglia as part of a coordinated neuroendocrine response. Future work defining the molecular states of ARC microglia and testing whether microglial manipulation is necessary or sufficient for PRP-induced metabolic effects will be important to clarify this mechanism.

Microglial morphology should not be interpreted as a binary indicator of inflammatory state, because microglia occupy diverse and dynamic functional phenotypes and soma area alone cannot reliably distinguish “activated” from “resting” cells^33^. Accordingly, the reduction in soma area observed here is best viewed as a morphological adaptation rather than evidence of generalized immunosuppression. This interpretation is consistent with our gene-expression data: *Tnf* increased, whereas *Il6*, *Il1b*, and *Nfkbia* were unchanged, arguing against a broad dampening of canonical inflammatory programs. Instead, the combined morphological and multidirectional transcriptional changes point to a more nuanced remodeling of the hypothalamic neuroimmune milieu. Defining the functional significance of this response will require additional measures of microglial state and activity, including branching complexity, phagocytic capacity, interactions with metabolically relevant neuronal populations, and cell-type-resolved transcriptional profiling.

At the pathway level, the transcriptomic findings further support the conclusion that central PRP engages multiple hypothalamic processes, affecting immune and metabolic signaling, extracellular matrix/cell-adhesion programs, and growth-related responses rather than uniformly suppressing inflammatory genes. Notably, altered *Lepr* and *Npr3* suggest potential links to neuroendocrine signaling, while changes in *Dcn*, *Spp1*, MHC class II–associated genes, *Bmp6*, *Bmp7*, and *Ptgdr* implicate tissue organization, immune communication, growth-factor signaling, and lipid mediator pathways. qPCR validation supported selected RNA-seq-associated changes. Although these data do not establish causality for any single pathway, they provide a molecular roadmap to prioritize candidate mediators and responsive hypothalamic cell populations for future mechanistic studies.

The broad transcriptional response observed in the hypothalamus is biologically plausible because PRP is a complex, multicomponent preparation rather than a single-factor intervention. Its growth factors, cytokines, chemokines, lipid mediators, and extracellular vesicles may act simultaneously on neurons, microglia, astrocytes, and vascular-associated cells, engaging distinct receptors and initiating both direct intracellular signaling and secondary paracrine responses. PRP-derived extracellular vesicles may further broaden and prolong these effects by transferring proteins, lipids, and nucleic acids between cells^34–36^. Such coordinated actions could account for the concurrent changes in immune communication, metabolic and neuroendocrine signaling, extracellular matrix and cell-adhesion programs, and growth-related pathways identified by RNA sequencing. This interpretation is supported by earlier work showing that PRP can influence neural repair, angiogenesis, axonal growth, and cultured central nervous system cell populations^16–20^. The present findings extend these observations by demonstrating that, even in the absence of focal CNS injury, central PRP can induce a broad hypothalamic transcriptional response in the context of chronic metabolic disease.

Notably, these interpretations should be viewed in light of several limitations. First, PRP composition was only partially characterized. Platelet enrichment was inferred from an independent validation cohort, whereas leukocyte carryover, platelet activation status, concentrations of soluble mediators, and extracellular vesicle content were not quantified in the administered preparations. In addition, sample freezing/thawing may have altered platelet integrity and the release profile of bioactive factors. Inclusion of vehicle-matched platelet-poor plasma and processing controls in future studies will be important to disentangle the relative contributions of platelets, residual plasma constituents, and handling conditions. Second, while intracerebroventricular delivery demonstrates CNS responsiveness, it does not identify the anatomical locus or the specific hypothalamic cell populations mediating the response. Finally, key endpoints (metabolic, histological, and transcriptomic readouts) were obtained in separate cohorts and at different time points, and the study was restricted to male mice, limiting inference about temporal coupling across outcomes and generalizability across sexes. Collectively, these constraints preclude attribution of the observed effects to any single PRP component or pathway and highlight priorities for more mechanistic, rigorously controlled neuroendocrine studies.

In conclusion, a single central PRP injection in DIO mice produced robust weight loss and coordinated changes in energy homeostasis, accompanied by reduced ARC microglial soma area and broad hypothalamic transcriptional changes. These findings extend PRP research from predominantly regenerative applications to central metabolic regulation and connect platelet biology with hypothalamic neuroimmune and metabolic physiology. This proof-of-concept phenotype provides a foundation for identifying the active PRP components, responsive brain cells, and neuroendocrine circuits that may inform future biological strategies for obesity and metabolic disease.

## Acknowledgments

The authors gratefully acknowledge the Biologic Resources Laboratory at the University of Illinois Chicago for assistance with mouse colony maintenance and the Metabolic Phenotyping Core at the University of Illinois Chicago for support with indirect calorimetry measurements.

This work was supported by the Key Research and Development Program of the Health Commission of Ningxia Hui Autonomous Region (Grant No. 2024-NWZD-B002).

## Author Contributions

PX and PL conceived and designed the study. XH and XYY performed the experiments and collected the data, with contributions from XHY, QX, MDM, MM, DJ, PX, and PL. XH and PL analyzed the data and prepared the figures. PX and PL drafted and revised the manuscript and supervised the study. XYY secured funding. All authors have read and approved the final manuscript.

