## Supplemental figures for "Central platelet-rich plasma administration promotes weight loss in diet-induced obese mice"

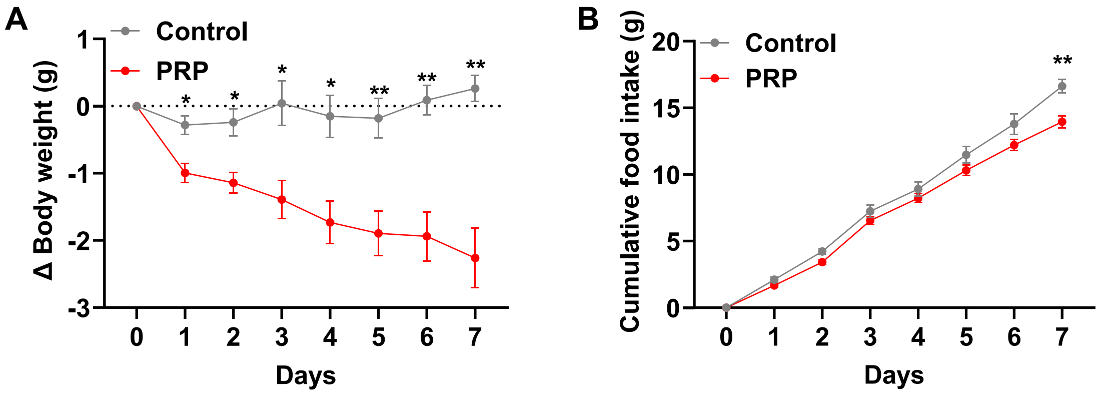


**Figure S1. Effects of central PRP on body weight and food intake in diet-induced obese mice.**

Diet-induced obese mice received a single stereotaxic ICV injection of PRP or aCSF directly into the lateral ventricle without prior ICV cannulation, in contrast to the cannula-based administration paradigm used in Figure 2B–C. Body weight change (A) and cumulative food intake (B) were monitored for 7 days after ICV administration. Data are presented as mean ± SEM; n = 10 mice per group. Statistical significance was analyzed by two-way ANOVA with Sidak's multiple-comparisons test (A, B). Significance thresholds: $*P < 0.05$, $**P < 0.01$.

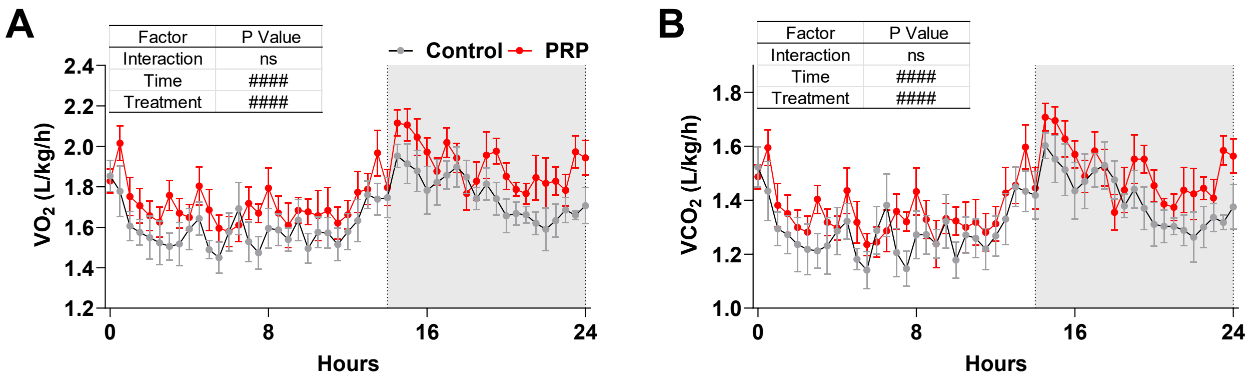


**Figure S2. Effects of central PRP on oxygen consumption and carbon dioxide production in diet-induced obese mice.**

**(A)** Oxygen consumption ($VO_2$) over the 24-h recording period.

**(B)** Carbon dioxide production ($VCO_2$) over the 24-h recording period.

Data are presented as mean ± SEM; n = 7 mice per group. Data were analyzed by two-way ANOVA (A, B). Significance thresholds: ####P < 0.0001.

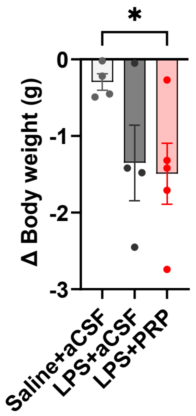


**Figure S3. Body weight change following LPS and central PRP treatment.**

Body weight change in Saline+aCSF, LPS+aCSF, and LPS+PRP groups.

Data are presented as mean ± SEM; n = 5 mice per group. Statistical significance was analyzed by ordinary one-way ANOVA followed by uncorrected Fisher's LSD multiple-comparisons test. Significance thresholds: $^*P < 0.05$.

**Table S1. Primer sequences used for quantitative PCR.**

| Gene | Forward primer (5′–3′) | Reverse primer (5′–3′) |
|---|---|---|
| *Gapdh* | AGGTCGGTGTGAACGGATTTG | TGTAGACCATGTAGTTGAGGTCA |
| *Il6* | GTGGCTAAGGACCAAGACCA | GGTTTGCCGAGTAGACCTCA |
| *Il1β* | GAAATGCCACCTTTTGACAGTG | TGGATGCTCTCATCAGGACAG |
| *Tnfα* | CATCTTCTCAAAACTCGAGTGACAA | TGGGAGTAGATAAGGTACAGCCC |
| *Nfkbia* | TGAAGGACGAGGAGTACGAGC | TGCAGGAACGAGTCTCCGT |
| *Lepr* | CAGCCCATGCTTGTTGTGAA | GTGCCATTGTTTGGCTGTCC |
| *Npr3* | TTGCACACGTCTGCCTACAA | ACACATGATCACCACTCGCT |
| *Ccn2* | AGAACTGTGTACGGAGCGTG | GTGCACCATCTTTGGCAGTG |
| *Dcn* | GCAAATACCCGGATTAAAAGGTCG | AATGGTCCAGCCCAAGAGAC |
| *Spp1* | GCTTGGGTTTGCAGTCTTCTG | TCTCATGGTCGTAGTTAGTCCCTC |
| *H2-Aa* | GTTCTGAAACACTGGGAACCTG | GATCGCAGGGCCTTGAATGATG |
| *H2-Ab1* | GTCACAGGAGTCAGAAAGGACC | GCAGTCAGGAATTCGGAGCA |
| *H2-Eb1* | AAGATGTTGAGTGGAGTTGGGG | CTGAAGTCCAGACTGTCCTTTCT |
| *Bmp6* | GCTTTGTGAACCTGGTGGAGTA | TCACCCTCAGGAATCTGGGATA |
| *Bmp7* | GAGCTTCGTCAACCTAGTGGAA | TAGATCCTGAATTCGGCTGCG |
| *Ptgds* | ACCATCCCCACTGTGAGTTTT | ACCAAACCCATCCACAGCAT |
| *Ptgdr* | GTTCGCCTTCCTGATGTCCT | AGGTGACGTGCCTTTGGTAG |
